# The role of the Angular Gyrus in the Elaboration of Specific and Categoric Memories

**DOI:** 10.64898/2026.07.17.739117

**Authors:** Alice Bush, Fiona Lancelotte, Ann-Kathrin Johnen, Greta Melega, Dingrong Guo, Cristian Lopez Saquisili, Elizabeth Jefferies, Jonathan CW Brooks, Louis Renoult

**Affiliations:** School of Psychology, University of East Anglia, Norwich, UK; School of Psychology, University of Sussex, UK; Department of Psychology, Birmingham City University, UK; Department of Neurology, Charité, Universitätsmedizin Berlin, Berlin, Germany; Department of Psychology and Cooperative Brain Imaging Centre, Goethe University Frankfurt, Frankfurt, Germany; Department of Psychology and Human Development, University of East London, London, UK; Department of Psychology, University of York, York, UK; Department of Psychology, University of Liverpool, Liverpool, UK

**Keywords:** Angular Gyrus, Default Mode Network, Autobiographical memory, Memory Details

## Abstract

This study investigated the role of the angular gyrus (AG) and its subregions in the elaboration of specific and categoric autobiographical memories (AMs). Using a cue-word fMRI paradigm, thirty-nine participants retrieved and elaborated on episodic (specific) and semantic (categoric) memories, rating the amount of detail of each recollection. Parametric analyses revealed that AG activity, particularly in posterior AG, was positively associated with the amount of detail retrieved, regardless of memory type. Specific memories elicited greater activation in right AG subregions compared to categoric memories. These findings support the AG’s involvement in both episodic and semantic memory retrieval and suggest functional differentiation between its subregions.

---

The distinction within declarative memory between episodic (memories of personally experienced unique events) and semantic memory (conceptual and factual knowledge) remains of central importance in cognitive neuroscience. In recent years, investigators have noted a striking similarity in their neural substrates, both engaging regions of the default mode network, also referred to as core recollection network or core network (DMN; Addis, 2018, Rugg & Vilberg, 2013, Addis et al., 2007; Binder & Desai, 2011; Renoult et al., 2019; Irish & Vatansever, 2020). Though often studied separately, both episodic and semantic information make-up the content of our autobiographical memories (AMs). For example, based on context or specific tasks demands, one may retrieve more specific AMs (e.g., the first time I climbed a route outdoors) or semantic or categoric AMs (g., hiking in the Alps every summer). AM studies reveal that brain regions like the hippocampus and precuneus are commonly found to show greater level of activity for specific AMs, while temporal lobe areas are thought to be key to support categoric retrieval (reviewed in Sheldon et al., 2019; St Jacques, 2012; Addis et al., 2016). However, most AM studies have focused on episodic memory, typically instructing their participants to retrieve specific AMs. DMN regions like the angular gyrus (AG) have been identified as key in both episodic (Rugg & King, 2018; Simons et al., 2022; Wagner et al., 2015) and semantic memory studies (Binder & Desai, 2011; Fernandino et al., 2016; 2022; Kuhnke et al., 2023) and AM studies offer a unique opportunity to clarify their functional role in declarative memory retrieval, provided that episodic and semantic conditions are included (e.g., Addis et al., 2004; Holland et al., 2011). In the present fMRI study, we used a paradigm in which participants were asked to retrieve specific or categoric memories in different trials to clarify the role of the AG and subregions in episodic and semantic retrieval, as well as of other key regions like the hippocampus and the precuneus commonly involved in AM retrieval.

Situated within the default mode network (DMN), the AG lies in the inferior parietal lobe (IPL), at the convergence of subsystems associated with the integration of multimodal perceptual cues and semantic features (Andrews-Hannah et al., 2010; Jefferies & Smallwood, 2025; Lanzoni et al., 2020; Murphy et al., 2019; Price et al., 2015; Yeo et al., 2011). This positioning has made it difficult to determine whether AG activity fluctuates with memory type (episodic versus semantic) or instead reflects domain general processes relating to experiential qualities such as memory vividness and richness. Lesion and TMS stimulation studies indicate that disruptions to IPL leads to impoverished autobiographical recollection, particularly in terms of retrieved detail (Berryhill et al., 2007; Berryhill, 2012; Bonnici et al., 2018). Functional neuroimaging studies have similarly linked AG activity to the amount and quality of retrieved information rather than to retrieval success alone (Rugg & King, 2018; Tibon et al., 2019; Yu et al., 2012). Parallel investigations on the role of AG in semantic cognition reported that it was modulated by the number of semantic features of concepts (Ferreira et al., 2015), by their concreteness or imageability (Graves et al., 2010; Binder et al., 2005; Sabsevitz et al., 2005) and that it showed greater activity for semantically congruent than incongruent stimuli (e.g., Kuhne et al., (2023). The involvement of the AG in the retrieval of semantic memory (Binder et al., 2009; Binder & Desai, 2011; Fernandino et al., 2016; 2022; Kuhnke et al., 2023) can be related to the proposal that recollection effects in the core recollection network may reflect reinstatement of conceptual processing, or may be related to the amount of information being retrieved during both semantic and episodic tasks (Renoult et al., 2019; Rugg & King, 2018; Rugg, 2022). To reconcile the functional role of the AG in semantic and episodic retrieval, the ‘dynamic buffer hypothesis’ proposes that the AG enables a temporary (online) multimodal buffering of internal and external information (Humphreys et al., 2021). This hypothesis considers the AG as a domain-general region, supporting its involvement in a wide array of cognitive functions.

A further relatively underexplored issue is whether AG subregions have different roles in memory retrieval. The AG is subdivided into two cytoarchitectonic subregions, anterior (PGa) and posterior (PGp) (Caspers et al., 2006; Seghier, 2013). These differ in their connectivity, with PGa being more strongly connected with frontal and cingulate regions, and PGp with the core recollection network, including hippocampus, medial prefrontal cortex, and precuneus (Uddin et al., 2010). Despite this, relatively few studies have examined memory-related processes in these subregions, and the method of subdivision has varied across studies (Bellana et al., 2023; Humphreys et al., 2022; Humphreys et al., 2024; Kunkhe et al., 2023). Existing evidence suggests that PGp is sensitive to both episodic recollection and access to prior knowledge (Bellana et al., 2023), while both PGa and PGp can be modulated by semantic demands (Kuhnke et al., 2023). However, anterior-posterior differentiation within ventral lateral parietal cortex has been described, with PGa preferentially connected to language and semantic processing systems, and more posterior regions linked to visual and attentional processing, while an intermediate posterior AG region (mid-PGp) shows strong links with the episodic recollection network (Humphreys et al., 2022).

This graded organisation within the AG aligns with connectivity parcellations of inferior parietal cortex, in which AG spans multiple DMN subdivisions (Andrews-Hannah et al., 2010; Jefferies & Smallwood, 2025; Yeo et al., 2011), providing a network-level framework for examining how cytoarchitectonic subregions map onto DMN subsystems. Recent work has linked these DMN subdivisions to partially distinct cognitive functions. The dorsomedial subsystem (including lateral and anterior temporal lobes, AG, and dorsomedial prefrontal cortex) shows the strongest overlap with semantic cognition (Andrews-Hannah et al., 2010; Jefferies & Smallwood, 2025). The medial-temporal subsystem (left posterior AG, lateral occipital cortex and medial temporal cortex) is associated with scene construction and contextual retrieval, whereas core DMN regions (posterior cingulate, AG) are engaged when cues and features are integrated (Andrews-Hanna et al., 2010; Shao et al., 2024; Jefferies & Smallwood, 2025). Consequently, in addition to examining cytoarchitectonic subdivisions (PGa and PGp), we assessed AG organisation using core, dorsomedial and medial-temporal DMN subdivisions. This complementary approach allowed us to test whether anterior-posterior effects observed across PGa and PGp were mirrored across DMN subsystems differing in their involvement in semantic cognition (dorsomedial), episodic retrieval (medial-temporal), and integrative memory processes (core DMN).

To clarify the role of the AG in episodic and semantic retrieval, and its sensitivity to the richness of information being retrieved, the present study compared the retrieval of specific versus categoric AMs in a cue word paradigm similar to the Autobiographical Memory Test (AMT; Williams & Broadbent, 2000). Participants first received a probe indicating whether they would have to think about a specific or categoric memory in the next trial. They were then presented a word cue (e.g., “Apple”) and asked to press a key when they had a word-related memory in mind (“Continue thinking about the details of the memory”). Due to our interest in the sensitivity of the AG to the richness of retrieved episodic and semantic content, we focused our analyses on the elaboration rather than the search phase (Daselaar et al., 2008; though a whole-trial model, collapsing across search and elaboration phases is reported in the supplementary section). We predicted that AG activity would be higher for both types of memory as compared to the control task, and we expected a positive relationship between activity in the left AG (specifically PGp) and number of details retrieved.

## Methods

### Participants

Forty-four healthy right-handed young adults, with no prior history of neurological or psychiatric impairment, participated in this study. Five participants’ data were removed from the analysis: two due to excessive movement during scanning (MCFLIRT mean displacement was greater than 3mm), and three due to not having completed the task per instructions (failing to press a key, indicating the retrieval of a memory and the start of the elaboration phase for both categoric and specific memories). Of the remaining thirty-nine participants, 26 were females, and their mean age = 23.38 years (ranging from 18 to 34, SD = 4.68).

### Stimuli

Cue words used in the fMRI AM task were 30 nouns selected from the Clark and Paivio (2004) extended norms. All were high in frequency (M = 1.55, SD = .42; Thorndike & Lorge, 1994), imageability (M = 5.77, SD = .40) and concreteness (M = 6.86, SD = .29) to increase the likelihood that an event could be generated by participants. For each participant fifteen words were randomly assigned to each condition (specific memory, categoric memory; see below). Participants were also presented fifteen letter arrays for a letter change control task (45 total).

### Procedure

Participants completed an autobiographical memory task, consisting of specific (episodic), categoric (semantic), and control trials, presented in alternating blocks; each block including five trials. Two different versions of the task were used, this was so all cue words were seen as both specific and categoric, with each participant seeing each word once. For memory trials, participants were cued to retrieve either a specific event or a repeated categoric experience associated with a cue word. Once a memory came to mind, participants pressed a button, marking the transition from a search (construction) phase to an elaboration phase during which they continued thinking about the memory. After elaboration, participants rated the amount of detail retrieved on a 4-point Likert scale (1 = no details; 4 = highly detailed). Control trials consisted of a letter change detection task. Participants were instructed to press a button when a letter change was detected, thereby matching motor demands of the memory trials. A practice task, including specific, categoric memories and control trials was administered prior to scanning (computer task using PsychoPy 2022.2.4).

### Image Acquisition

Imaging data were acquired using a 3 T Siemens Prisma MRI scanner with a 32-channel head coil. Structural images were acquired using sagittal T1-weighted MPRAGE pulse sequence, TE/TI/TR = 2.17/900/2200ms, flip angle = 9°, GRAPPA acceleration factor = 2, voxel size = 1.0 × 1.0 × 1.0mm, and bandwidth = 210Hz/Px. Subsequently, functional data were acquired in a transversal orientation, with an echo-planar imaging (EPI) sequence: TR/TE = 2000/30ms, flip angle 76°, multi-band acceleration factor = 3, resolution 3 × 3 × 3mm, bandwidth = 2604 Hz/Px, echo spacing of 0.49ms and with interleaved multi-sliced mode, with 51 slices. The memory task had a duration of 708 measurements (i.e., 23m36s). At the end, a field mapping sequence was acquired in a transversal orientation, with the following sequence parameters TE1/TE2/TR = 4.92/7.38/520ms, flip angle 60°, resolution 3 × 3 × 3mm, bandwidth = 596 Hz/Px and with interleaved multi-slice mode, with 49 slices.

### fMRI Analysis

Functional images were pre-processed using FEAT (FMRI Expert Analysis Tool) version 6.00, part of FSL (FMRIB’s Software Library, www.fmrib.ox.ac.uk/fsl). The following preprocessing steps were applied; motion correction using MCFLIRT (Jenkinson et al., 2002); nonbrain removal using BET (Smith, 2002); spatial smoothing using a Gaussian kernel of FWHM 5mm; grand-mean intensity normalisation of the entire 4D dataset by a single multiplicative factor; highpass temporal filtering (Gaussian-weighted least-squares straight line fitting, with sigma=45.0s). Motion correction and fieldmap unwarping were performed in a single step to minimise the amount of smoothing arising from multiple interpolation steps. Registration of motion-corrected, unwarped, functional imaging data to a standard was performed using Boundary Based Registration (BBR, Greve & Fischl, 2009) and 6 degrees of freedom (dof) linear registration to each subject’s own skull stripped T1-weighted structural, followed by 12 dof affine registration using FLIRT (Jenkinson et al., 2002) and non-linear registration to the MNI152 T1-weighted 2mm resolution template brain (Andersson et al., 2007a; 2007b). First-level analyses were conducted for each participant’s data in acquired space (i.e., not normalized to MNI) using FEAT in FSL. Time-series statistical analysis was carried out using FILM with local autocorrelation correction (Woolrich et al., 2001). Simple main effects were estimated by creating difference contrast images between conditions at the first level. Three separate models were run:

1. Whole-trial model, collapsing across search and elaboration phases to characterise overall task-related activity associated with autobiographical memory retrieval.
2. Elaboration model, elaboration phases using the button-press timing, enabling isolation of activity during memory elaboration.
3. Parametric model, including trial-by-trial self-reported detail ratings as a parametric regressor during the elaboration phase.

All models included regressors of to account for key presses. For all regressors, a temporal derivative was included to help account for slight differences in the timing of BOLD responses. Group level analyses were conducted using mixed effects modelling (FLAME 1+2). Whole brain inference used cluster corrected thresholds (Z > 3.1, p < .05 FWE). For region of interest (ROI) analyses, non-parametric permutation testing was conducted using RANDOMISE, and significance assessed with threshold free cluster enhancement (corrected P<.05 FWE). Whole brain results from the whole trial model are reported in the Supplementary Materials and serve to validate task engagement and robustness of analyses. In line with the theoretical aims of the study, inferential analyses in the main manuscript focus on the elaboration and parametric detail models.

### Mask Selection

Cytoarchitectonically defined masks for PGa and PGp were derived from the Juelich Histological Atlas (Amunts et al., 2020) available in FSLeyes and analysed separately for left and right hemispheres. Additional masks included the precuneus and hippocampus, selected using the probabilistic Harvard–Oxford atlases. All masks were thresholded at P(region)>0.4 (i.e. greater than 40% probability of voxels belonging to selected region), consistent with previous studies (Bellana et al., 2023). To assess whether angular gyrus effects observed with cytoarchitectonic masks generalised to a network-level functional parcellation, additional masks were defined using the Yeo et al. (2011) 17 network. Functional parcels corresponding to DMN subdivisions: core, dorsomedial and medial-temporal were extracted separately for the left and right hemispheres from the Schaefer–Yeo 400 parcel MNI152 atlas (Schaefer et al., 2018). These subdivisions were selected based on previous work associating the dorsomedial subsystem with semantic cognition, the medial-temporal subsystem with episodic memory, and core DMN regions with integration of cues and retrieval of meanings from memory (Jefferies & Smallwood, 2025). To constrain these network parcels to inferior parietal cortex, all Jülich histological masks corresponding to inferior parietal lobule (IPL) regions were thresholded at P(region)>0.4 and summed to create hemisphere-specific “inferior parietalIPL40” masks. Each of the Yeo DMN parcels was then mapped onto the corresponding left or right IPL40 mask, yielding functionally defined masks within the IPL for each DMN subdivision. The dorsomedial DMN subdivision was not represented within the right IPL mask at the chosen threshold and was therefore included for left-hemisphere analyses only. Subsequently, ROI analysis was performed with the FSL tool FEATQUERY. The prespecified masks were transformed into the acquisition (i.e. functional) space for each subject, and percentage BOLD signal versus baseline was extracted for the contrasts of interest (e.g. elaboration phase for categorical memory trial).

## Results

### Behavioural Results

Firstly, the average number of responses to word cues in the memory task was calculated for each condition separately. Trials were retained only if participants pressed a button when they thought of a memory (either specific or categoric), as they were instructed to not press a button if they could not think of a memory. For specific memories, the number of trials ranged from 7 to 15 (the maximum possible), with an average of 14.3 (SD = 1.4) across participants. For categoric memories, the number of trials ranged from 9 to 15, with an average of 14.4 (SD = 1.3).

The duration of the search phase was compared between specific and categoric trials using a Wilcoxon signed-rank test. Results indicated no significant difference between categoric (M = 4.71, SD = 2.00) and specific (M = 5.12, SD = 2.05) search durations, *W*(39) = 269.50, *p* = .094, 95% CI [-0.82, 0.02] (all durations in seconds). For the elaboration phase, a Wilcoxon signed-rank test similarly revealed no significant difference between categoric (M = 15.29, SD = 2.00) and specific (M = 14.88, SD = 2.05) elaboration durations, *W*(39) = 511.50, *p* = .091, 95% CI [-0.02, 0.82]. Finally, Wilcoxon signed-rank tests compared search and elaboration phase durations, for specific and categoric trials. In both cases, search times were significantly shorter than elaboration times categoric: *W*(39) = 3.00, *p* < .001, mean absolute difference = 11.21, 95% CI [9.26, 11.08]; specific: *W*(39) = 1.00, *p* < .001, mean absolute difference = 10.56, 95% CI [9.26, 11.08]. Due to the significant differences between the durations of the search and elaboration phases, it was deemed inappropriate to directly contrast these components in the subsequent fMRI analyses. Self-reported detail scores were compared between memory conditions (categoric versus specific). A One-Way repeated measures ANOVA showed there was no significant difference in detail ratings between categoric (M = 2.76, SD = 0.30), and specific (M = 2.87, SD = 0.36) memories, F(1,38) = 3.62, p = 0.065.

### Whole Brain Analyses

Whole-brain analyses for categoric and specific memory retrieval during elaboration revealed recruitment of key regions associated with the core recollection network (Rugg and Vilberg, 2013; Figure 3), namely the left medial prefrontal cortex, angular gyrus and parahippocampal gyrus; posterior cingulate/retrosplenial cortex, lateral temporal cortex; bilateral precuneus and hippocampus; left Broca’s areas, cerebellum, and bilateral subregions of the visual cortex.

**Figure 1.**
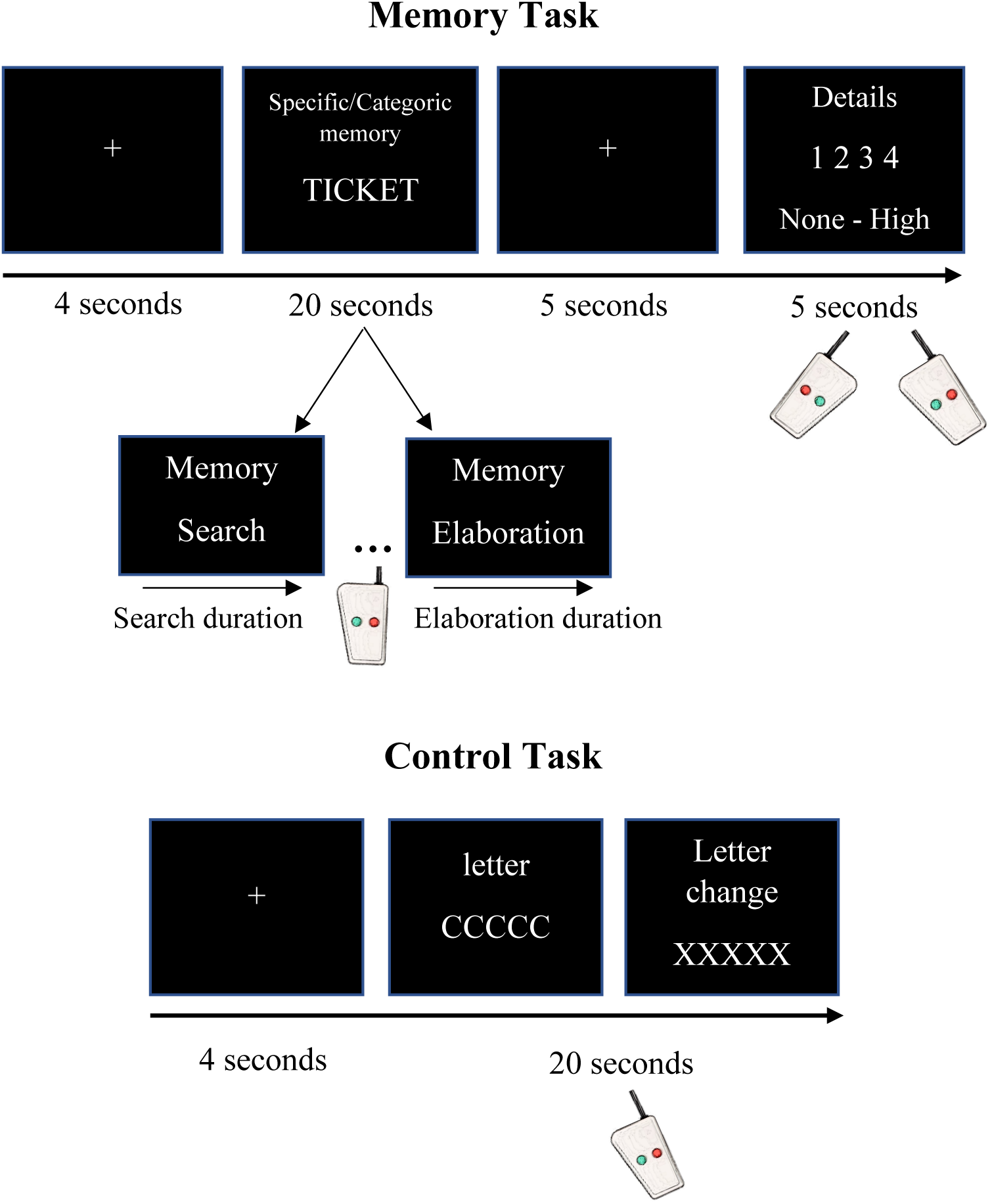
Task procedure. Participants were presented with a cue word and instructed to think about a specific or categoric memory (search phase). Once they had a memory in mind, they pressed a button, marking the onset of the elaboration phase. Participants were then asked to report how detailed their memory was on a scale of 1-4 (by pressing one of four buttons). The control task consisted of a letter change detection task, where participants pressed a button once they saw a change in the strings of letters being presented. Both memory (search + elaboration) and control tasks lasted 20 seconds.

**Figure 2.**
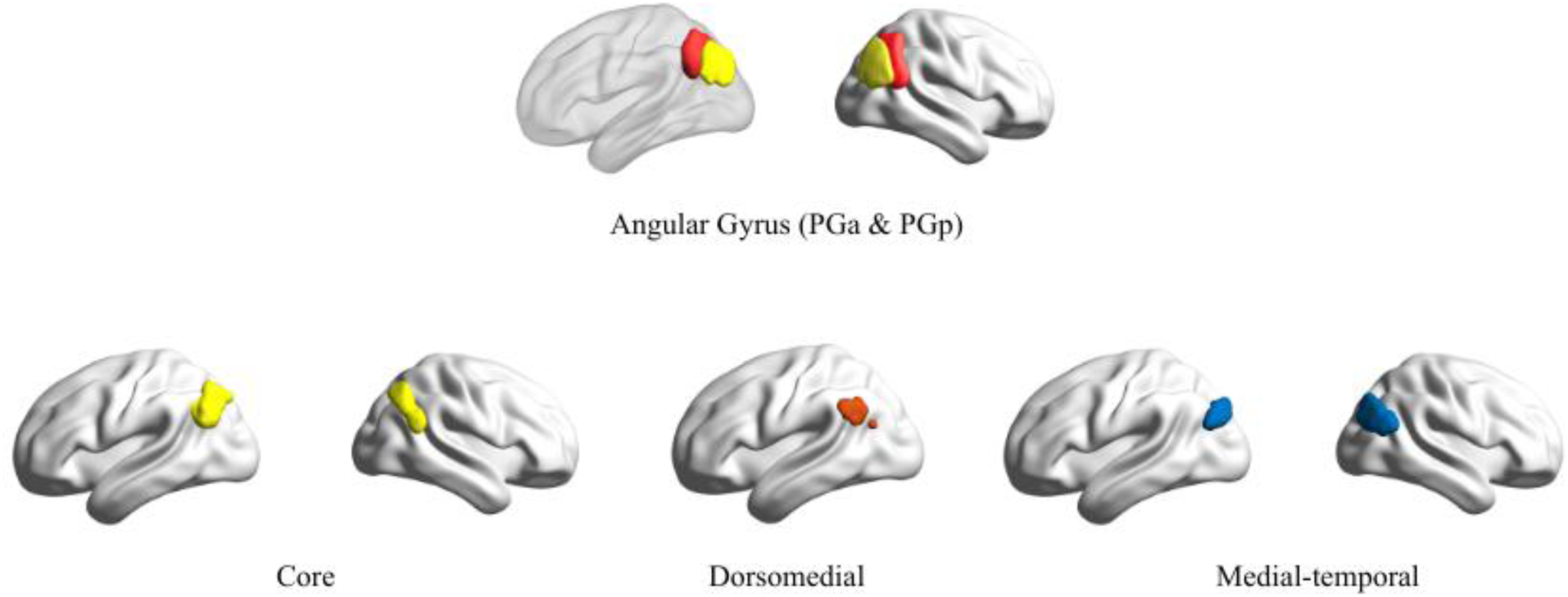
Masks derived from the probabilistic Jülich Histological Atlas (PGa in red and PGp, in yellow), and Yeo 17-network parcellation (Core, Dorsomedial, Medial-temporal), all thresholded at P(region)>0.4 displayed in the space of the MNI standard brain atlas.

**Figure 3.**
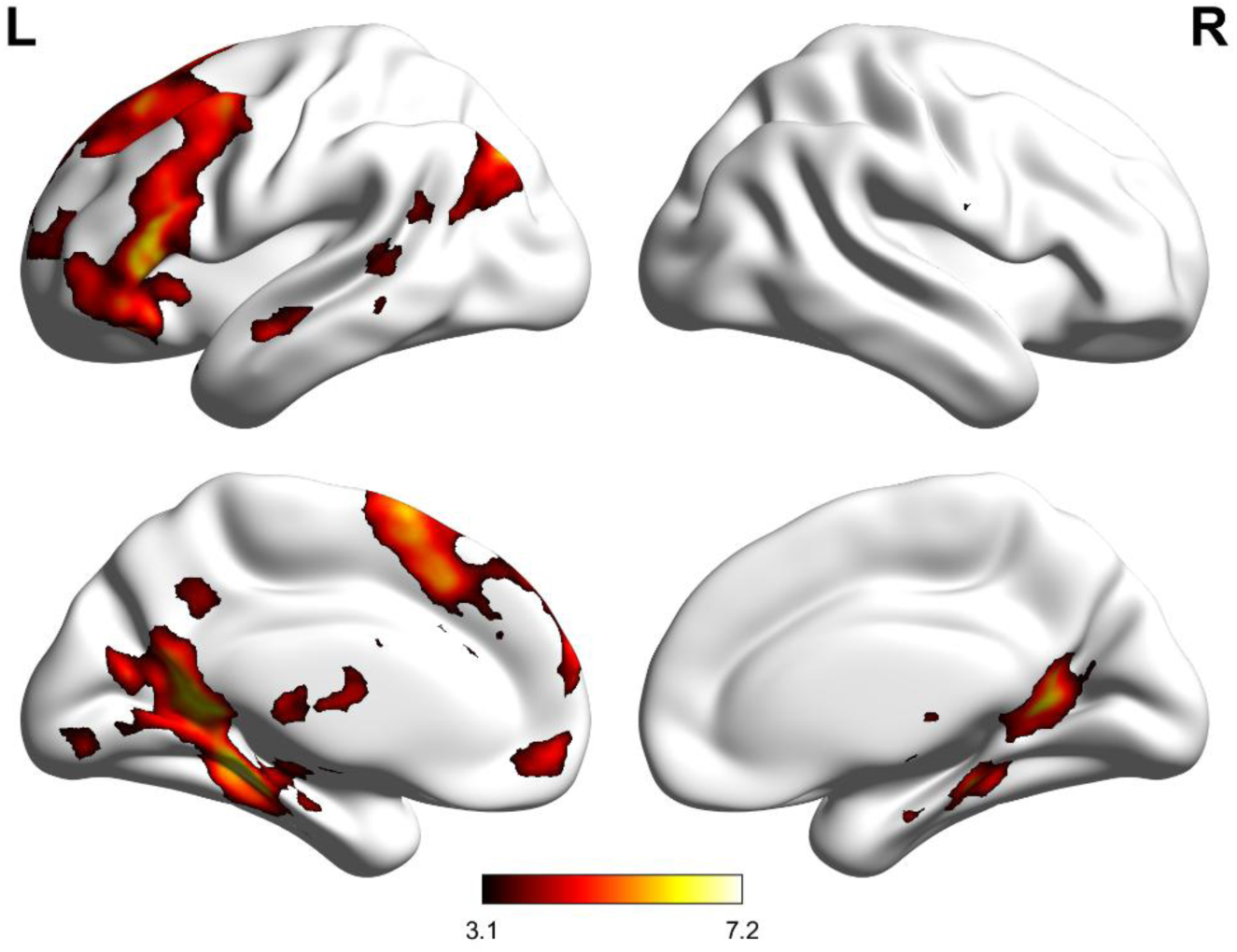
Activity for whole brain analysis of categoric + specific elaboration, versus the control task. Regions include: cerebellum; Frontal Pole, Middle Frontal Gyrus; Thalamus; Middle Temporal Gyrus; Precuneous; Parahippocampal Gyrus; Cingulate Gyrus; Hippocampus; Angular Gyrus; Inferior Parietal Lobule.

### Region of Interest Analyses

In line with our hypotheses, we wanted to investigate whether the AG is sensitive to the amount of detail retrieved during memory elaboration and whether its subregions would show different involvement.

Masked analyses were conducted to assess differences between specific and categoric retrieval. No masked analyses reached significance for the categoric > specific contrast. However, greater activity was seen during specific elaboration, compared to categoric elaboration for right PGa and PGp and precuneus. When assessing activity with the parametric regressor of details, a significant positive relationship was observed in left PGp and bilateral hippocampus during categoric retrieval, and bilateral PGp, right PGa, bilateral hippocampus and precuneus during retrieval of specific memories. Finally, when assessing whether activity based on the level of details recalled was differentially recruited for specific, compared to categoric memories, a stronger positive relationship was seen for the right AG (including PGa and PGp), left hippocampus and precuneus (See Supplementary Materials for all masked analyses).

**PGa and PGp subdivision.** To investigate the role of AG subregions in the retrieval of specific and categoric memories, we conducted ROI analyses for PGa and PGp during the elaboration phase. A 2×2 repeated measures ANOVA was conducted with factors of subregion (PGa & PGp) and memory type (categoric & specific) during elaboration (left hemisphere only). A significant difference of BOLD activity between subregions was found, F(1,38) = 7.408, *p* = 0.010 η²p = 0.163, but not between types of memory: F(1,38) = 0.0129, *p* = 0.910. There was also no significant interaction between main effects of region and memory type.

The same analysis was conducted for the right hemisphere (Subregion x Memory type). A significant difference of BOLD activity was found again between PGa and PGp F(1,38) = 67.96, *p* < .001, η²p = 0.641, but not between types of memory: F(1,38) = 0.078, *p* = 0.087. A significant interaction was found between subregion and memory type: F(1,38) = 10.62, *p* = 0.002, η²p = 0.218. Post-hoc Bonferroni corrected tests revealed greater activity for specific, compared to categoric memories in PGa (t(38) = 2.841, *p* = 0.043), but not PGp. Figure 4 below shows bilateral main effects of subregion (PGp > PGa), and subregion x memory type interaction for the right hemisphere (specific > categoric, in PGa).

**Figure 4.**
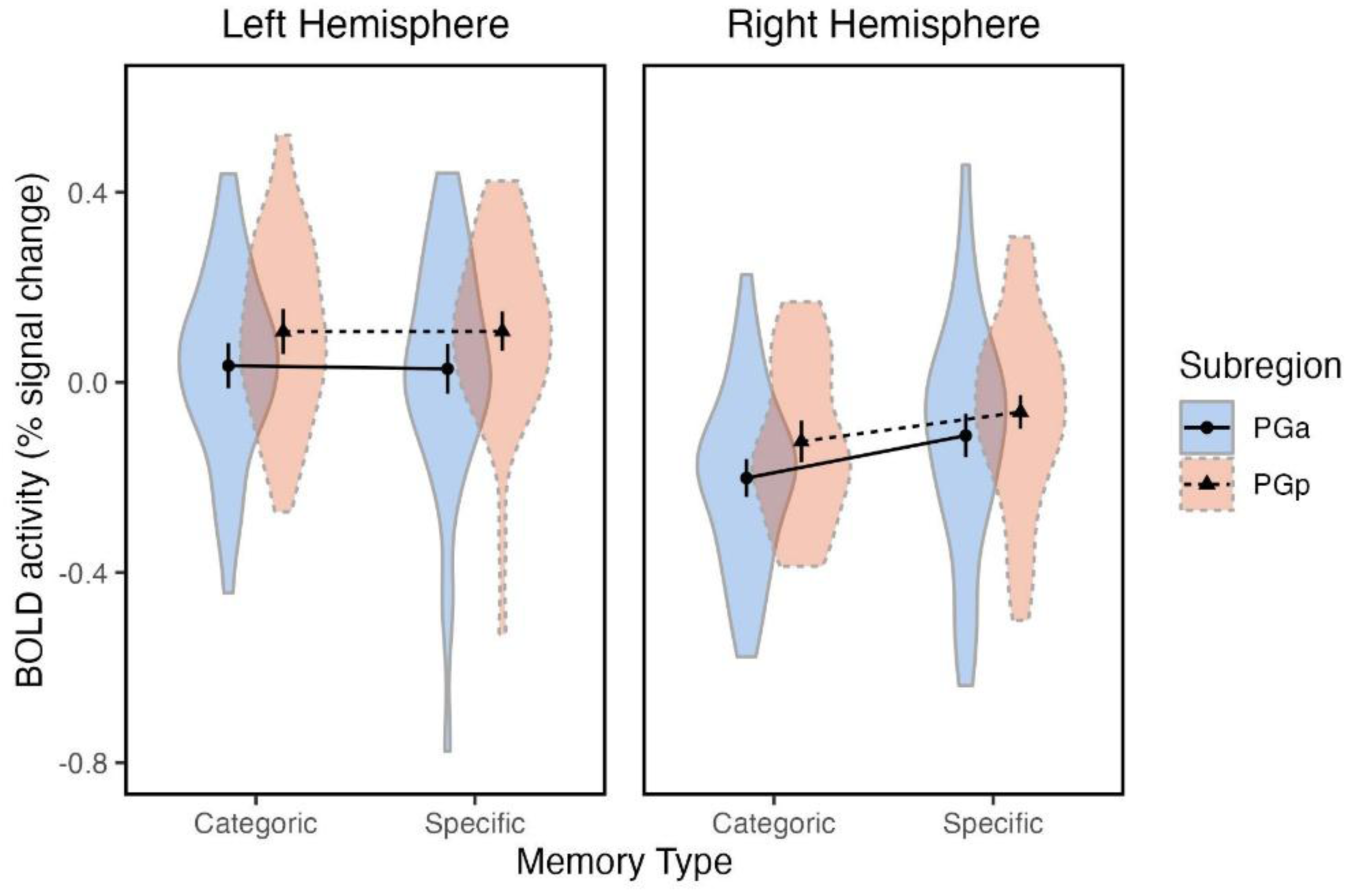
Mean BOLD signal (± SEM) during elaboration phase, for categoric and specific memory across PGa and PGp in the left and right hemispheres.

The same analysis was also conducted for the parametric analysis (elaboration with each trial weighted by subjects’ details rating). Similarly to the elaboration model above, the 2×2 repeated measures ANOVA showed a significant difference of BOLD activity in the left hemisphere between subregions, F(1,38) = 13.284, *p* < .001 η²p = 0.259. However, no significant differences between memory types F(1,38) = 0.291, *p* = 0.592 and no significant interaction subregion and memory type was found: F(1,38) = 0.007, *p* = 0.557.

When analysing BOLD activity in the right hemisphere, a significant difference was found again between subregions F(1,38) = 8.799 *p* = .005, η²p = 0.188. Unlike the elaboration model, a significant main effect of memory type was also found F(1,38) = 5.225, *p* = 0.028, η²p = 0.121. However, there was no significant interaction between subregion x memory type F(1,38) = 0.007, *p* = 0.935. Figure 5 shows bilateral main effects of subregion (PGp > PGa) and right hemisphere main effect of memory type (specific > categoric).

**Figure 5.**
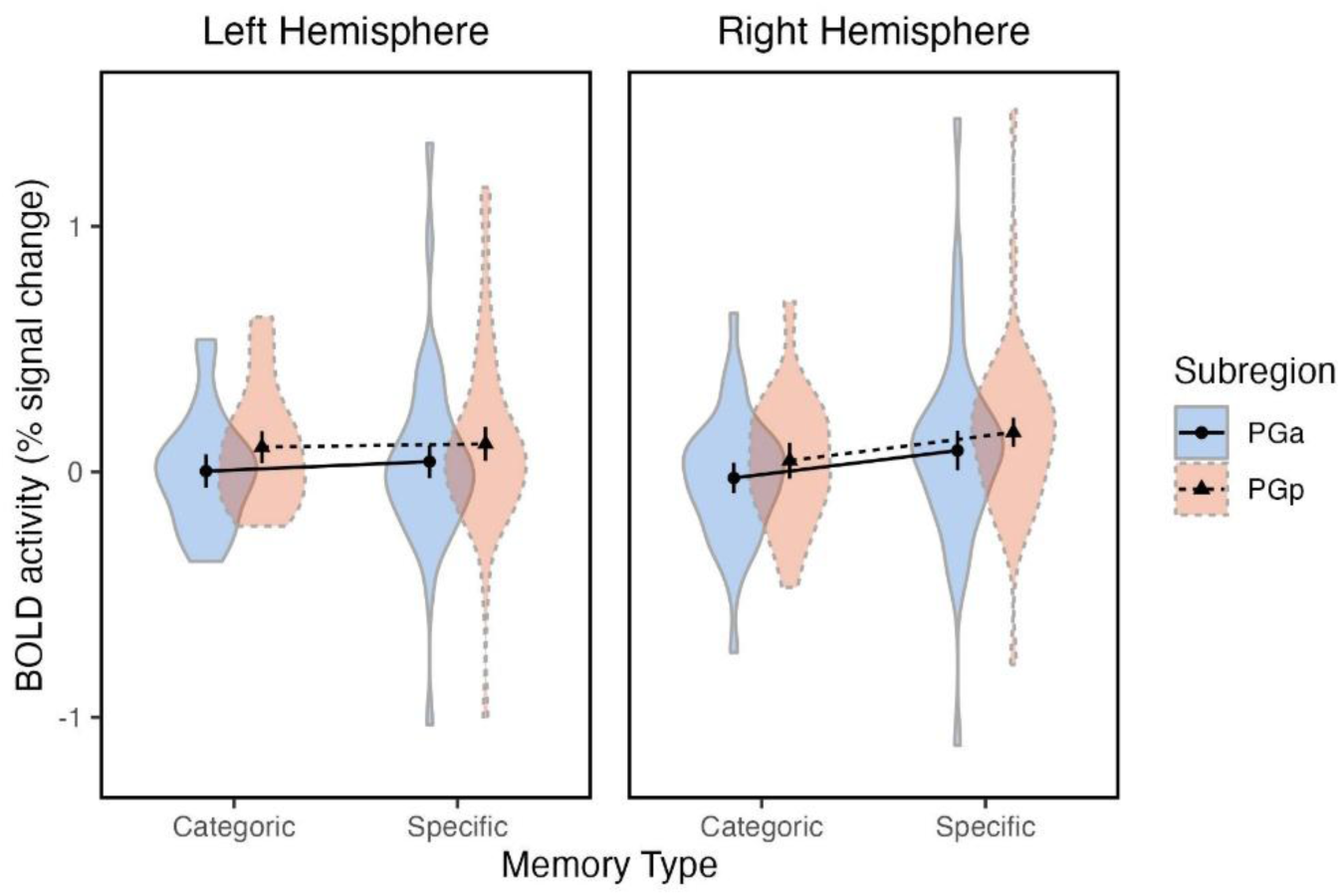
Mean parametric BOLD signal (± SEM) for elaboration phase with details as a regressor. 2×2×2 repeated measures ANOVA with factors of memory type (categoric and specific memory) across left and right AG subregions (PGa and PGp).

### DMN Subdivision within IPL: Core, Dorsomedial, Medial-temporal

Mask selection was created for core, dorsomedial and medial-temporal DMN subdivisions corresponding to IPL ( P(region)>0.4). Dorsomedial subdivision was not represented within the right IPL mask at the chosen threshold and was therefore included for left-hemisphere analyses only. A 3×2 repeated measures ANOVA for the left hemisphere with main factors region (core, dorsomedial, medial-temporal), and memory type (categoric & specific) during elaboration was conducted. A significant main effect of region was found (F(1,38) = 10.944, p < .001, np2 = 0.224), but not of memory types (F(1,38) = 0.488, *p* = 0.489). There was a significant interaction between region x memory type (F = 3.573, p = 0.033, np2 = 0.086). Post hoc analyses revealed significantly more BOLD activity in the DMN/IPL core and medial-temporal subregions, compared to dorsomedial (t(38) = 6.190, *p* < .001; t(38) = 3.515, *p* = 0.017, respectively) during categoric retrieval. There was also a significant difference between core > dorsomedial activity during specific retrieval (t(38) = 3.137, *p* = 0.049), but no significant difference between medial-temporal and dorsomedial activity during specific memory retrieval or medial-temporal and core differences, for either specific or categoric memory. Figure 6 shows left hemisphere main effects of subregions (core & medial-temporal > dorsomedial) and significant interaction (core specific > dorsomedial specific; core and medial-temporal categoric > dorsomedial categoric).

**Figure 6.**
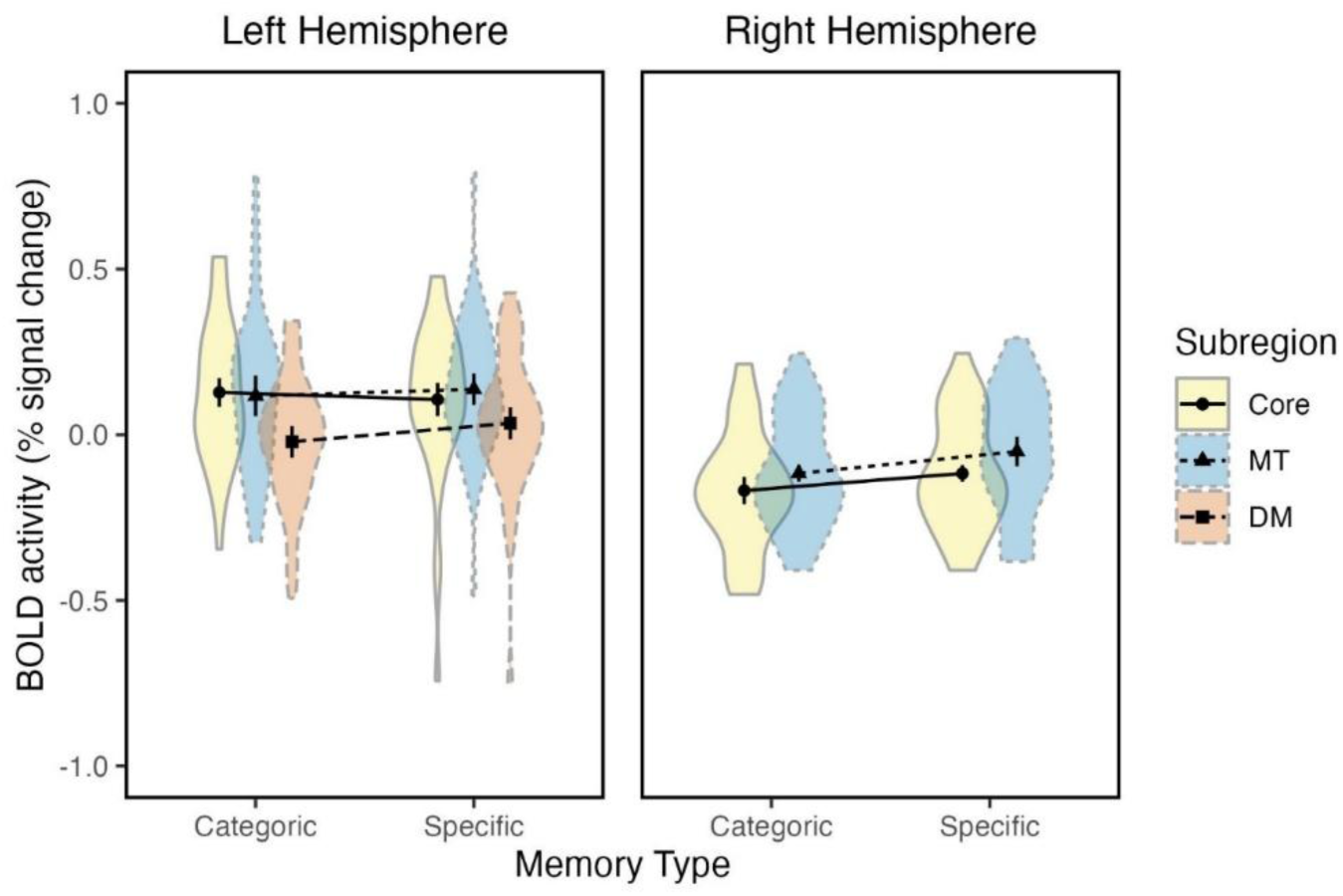
Mean parametric BOLD signal (± SEM) for elaboration phase. 3×2 repeated measures ANOVA with factors of memory type (categoric and specific memory) for left hemisphere DMN (Yeo 17) subregions (core, dorsomedial and medial-temporal) and 2×2 repeated measures ANOVA with factors of memory type (categoric and specific memory) for right hemisphere DMN (Yeo 17) subregions (core, and medial-temporal).

For the right hemisphere, a 2×2 repeated measures ANOVA with main factors region (core and medial-temporal), and memory type (categoric & specific) during elaboration was performed. A significant main effect of region was found (F (1,38) = 15.710, p < .001, np2 = 0.292), as well as a main effect of memory type (F (1,38) = 15.710, p < .001, np2 = 0.292). However, no significant interaction between region x memory type was observed (F(1,38) = 0.107, *p* = 0.745). Figure 6 shows main effects of subregion (medial-temporal > core) and main effects of memory type (specific > categoric).

The same analyses were conducted for the parametric model with details as the regressor weighting. For the left hemisphere we conducted a 3×2 repeated measures ANOVA with factors region (core, medial-temporal and dorsomedial), and memory type (categoric & specific). A significant main effect of region was found (F (1,38) = 8.444, p < .001, np2 = 0.182). No significant main effect of memory type or interaction (memory type x region) was found (F (1,38) = 0.396, p = 0.553; F (1,38) = 1.926, p = 0.153, respectively). Figure 7 shows main effects of subregion (medial-temporal > core & dorsomedial).

**Figure 7.**
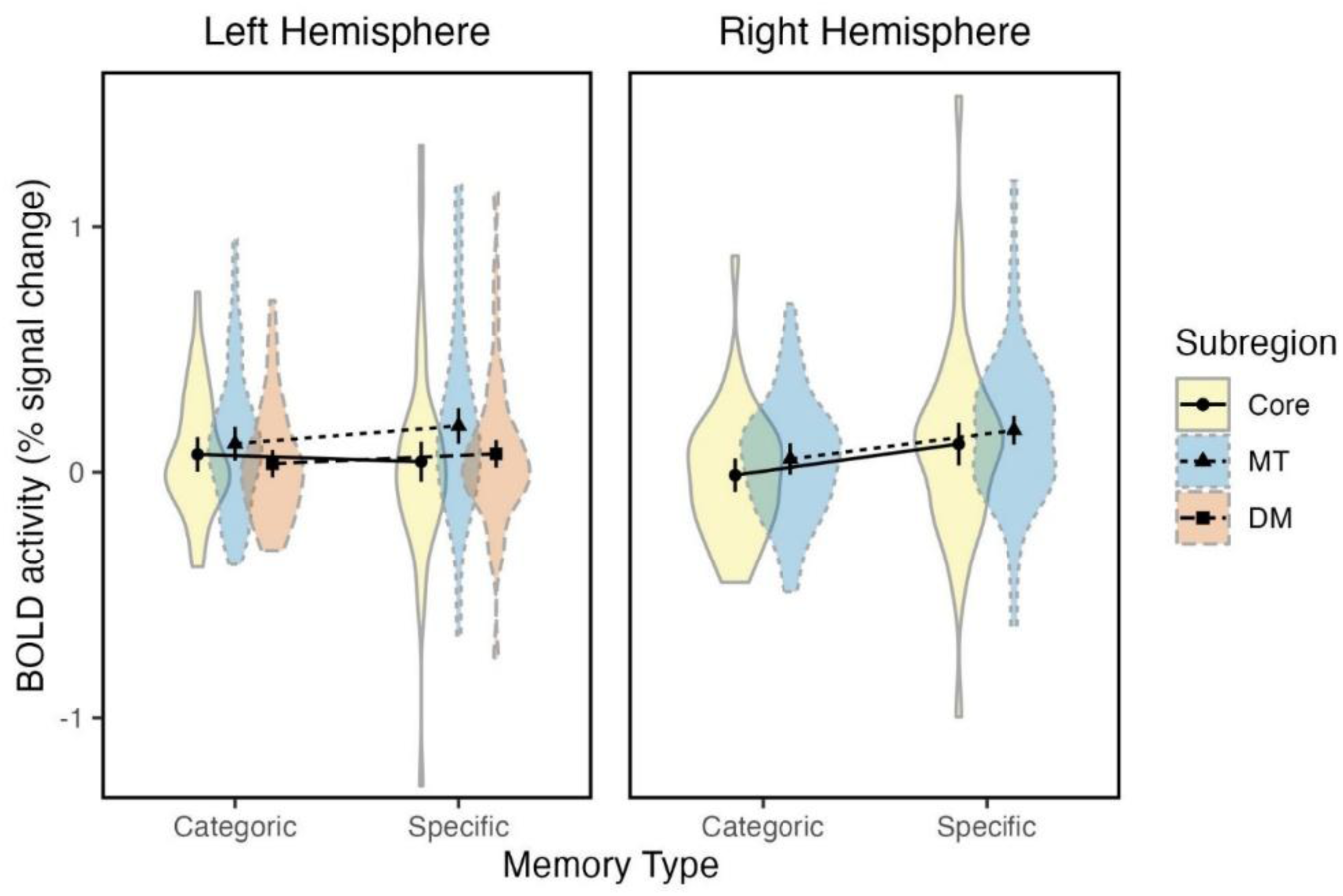
Mean parametric BOLD signal (± SEM) for elaboration phase, with details as a regressor. 3×2 repeated measures ANOVA with factors of memory type (categoric and specific memory) for left hemisphere Yeo-Julich masks (core, dorsomedial and medial-temporal) and 2×2 repeated measures ANOVA with factors of memory type (categoric and specific memory) for right hemisphere Yeo-Julich masks (core and medial-temporal).

For the right hemisphere, a 2×2 repeated measures ANOVA with factors region (core and medial-temporal), and memory type (categoric & specific) revealed a significant main effect of region (F = 5.478, p = .025, np2 = 0.126), as well as a main effect of memory type (F = 6.236, p = .017, np2 = 0.141). No significant interaction between region x memory type was observed (F(1,38) = 0.0387, *p* = 0.845). Figure 7 shows main effects of subregion (medial-temporal > core) and memory type (specific > categoric).

## Discussion

This study sought to clarify the functional organisation of the AG during AM retrieval and compare activity across episodic (specific) and semantic (categoric) retrieval. Our whole brain analysis revealed activation of the core recollection network (Rugg & Vilberg, 2013). Across analyses, left AG, particularly posterior subregion PGp, and DMN/IPL medial-temporal subregions were engaged during both specific and categoric memory retrieval and scaled with the richness of retrieved content. However, significantly higher BOLD activity was seen for specific than categoric retrieval in the right hemisphere. These findings support the view that left AG contributions to AM reflect integration and maintenance of internally generated representations, not exclusive to episodic or semantic memory. Finally, analyses for hippocampus and precuneus revealed bilateral hippocampal activity during elaboration, with details as a parametric regressor, irrespective of memory type (see Supplementary Materials).

### Comparing specific and categoric retrieval

Functional neuroimaging studies have demonstrated engagement of inferior parietal regions, including the AG, during perceptually decoupled states and during tasks requiring the integration of conceptual content across contexts (Murphy et al., 2018; Murphy et al., 2019). At the same time, other studies have reported apparent dissociations within the AG linked to task demands that emphasise either episodic recollection or semantic retrieval (Humphreys et al., 2022; Humphreys et al., 2024). Crucially, most previous work has tested episodic and semantic memory in separate paradigms, using different stimulus types and response demands. By prompting specific and categoric AM retrieval, the present study is distinct in examining a task context in which both episodic and semantic memory can be assessed within the same study.

By showing modulation of left AG by retrieval richness across both memory types, the current findings suggest that at least some dissociations previously attributed to memory category may instead reflect differences in experiential quality. This interpretation is consistent with lesion and stimulation evidence indicating that damage to inferior parietal cortex results in memories that are less vivid and less detailed, rather than categorically impaired (Berryhill et al., 2007; Berryhill, 2012; Bonnici et al., 2018), and with integrative accounts emphasising AG involvement in buffering multimodal content during recollection (Humphreys et al., 2021). However, there seems to be some differentiation of memory type within right hemisphere AG/IPL, with greater activity seen for specific, compared to categoric retrieval across ROIs.

### Subregional organisation of the angular gyrus

A central aim of the present study was also to clarify whether anterior and posterior AG subregions support distinct types of memory, or whether their recruitment is instead driven by experiential properties of retrieval. The present findings largely favour the latter interpretation. Behavioural results showed that equivalent number of details were reported by participants for specific and categoric memories. ROI analyses demonstrated greater activity for left PGp than left PGa, as well as for medial-temporal, than core and dorsomedial subregions of the DMN, irrespective of memory type.

Similar results were observed for right hemisphere ROIs, though, greater BOLD activity was seen for specific compared to categoric retrieval across PGa/PGp and DMN/IPL subdivisions. Prior research on episodic memory has commonly focused on left AG, rather than right or bilateral ROIs (Bellana et al., 2023; Holland et al., 2011; Humphreys et al., 2024) and left AG effects have been interpreted as reflecting the dependence of episodic memory on conceptual processing (Rugg & King, 2018). However, right hemisphere AG effects have also been reported in episodic memory experiments (e.g., Addis et al., 2004; Kwok et al., 2012; Flegal et al., 2014). In Addis et al., (2004) where a similar design as in the present study was used, specific and categoric memories both activated bilateral AG and hippocampus, compared to a control condition. The present findings suggest that left and right AG may be sensitive to distinct aspects of autobiographical retrieval. Whereas left AG activity scaled with retrieved detail across both memory types, right AG showed consistently greater activity for specific than categoric memories. One possible interpretation is that right AG is particularly sensitive to the retrieval of a unique autobiographical episode situated in a specific spatiotemporal context. This is consistent with previous studies linking right AG to episodic recollection and retrieval of temporal order information (Flegal et al., 2014; Kwok et al., 2012) and may explain why specific memories elicited greater activity despite equivalent subjective detail ratings.

An important implication of these results is that functional differentiation within the left AG may be better explained by the phenomenology of memory retrieval than by memory type per se. Cytoarchitectonic and connectivity studies have shown that anterior and posterior AG differ systematically in their long-range connections: anterior PGa is more strongly connected with frontal and temporal regions implicated in language and semantic control, whereas posterior PGp is more tightly coupled with the DMN and core recollection network including hippocampus, medial prefrontal cortex, and precuneus (Uddin et al., 2010; Humphreys et al., 2022; Seghier, 2013). The present findings extend this account by demonstrating that left posterior AG activity scales with retrieval richness not only during episodically focused retrieval but also during categoric autobiographical remembering. Such a pattern is more consistent with accounts proposing that AG operates as a domain-general buffer for internally generated representations (Humphreys et al., 2021), supporting the maintenance and integration of multimodal information during memory elaboration. This interpretation is also consistent with recent evidence that DMN subsystems show partially distinct response profiles while converging on common heteromodal regions during demanding retrieval processes (Souter et al., 2024), suggesting that functional differentiation within AG may emerge from its position across interacting DMN components rather than reflecting discrete memory systems. Thus, the network-level organisation of AG broadly mirrors the anterior-posterior gradients identified in cytoarchitectonic and connectivity studies. Importantly, greater activity for posterior subregions was observed across both cytoarchitectonic and network-level analyses. Within the Yeo-derived DMN parcellation (Yeo et al., 2011), the strongest responses were observed in the core and medial-temporal subdivisions, whereas dorsomedial regions showed comparatively weaker activation.

## Conclusion

In summary, the present findings demonstrate that AG involvement in AM appears to be largely driven by integrative and experiential demands, particularly within left posterior AG. However, right AG may retain greater sensitivity to the retrieval of specific autobiographical episodes. By combining cytoarchitectonic and network-level analyses within a paradigm that requires coordinated episodic and semantic processing, this study shows that left posterior AG consistently scales with retrieval richness across autobiographical memory types. The convergence of effects across PGa/PGp and DMN subdivisions highlights the importance of considering both local cortical architecture and large-scale network embedding when interpreting parietal contributions to memory. More broadly, these results help reconcile previously divergent accounts of AG function and support models in which posterior AG operates as a domain-general buffer for the elaboration of rich internal representations during memory retrieval.

## Acknowledgements

This project was funded by grant MR/S011463/1 from the Medical Research Council (MRC): “The neural correlates of personal semantic memory across the life course”.

