## Supplementary Material for "The role of the Angular Gyrus in the Elaboration of Specific and Categoric Memories"

**Yeo17 Subdivision Legend**

| Yeo et al., (2011) naming | Andrews-Hannah et al., (2010) naming |
|---|---|
| 15 | Medial-temporal |
| 16 | Core |
| 17 | Dorsomedial |

**Whole-Brain Analyses**

***Whole-Block Model***. Contrasts from the 'whole-block' model assessed the main effect of memory condition and demonstrated higher activation during specific, compared to categoric retrieval (Figure 1, Table 1) in the right paracingulate and cingulate gyri, middle temporal gyrus, supramarginal gyrus, and precuneus, the inferior parietal lobe (namely right PGa, left PF, and bilateral PFm), left cerebellum, and bilateral frontal regions, such as right Broca's area, inferior frontal gyrus (IFGT), frontal orbital cortex (OFC) and left frontal operculum cortex (FO), middle frontal gyrus and insular cortex (INS). No clusters met thresholding parameters of Z>3.1 and corrected significance level of p< .05 for the reverse contrast (categoric > specific).

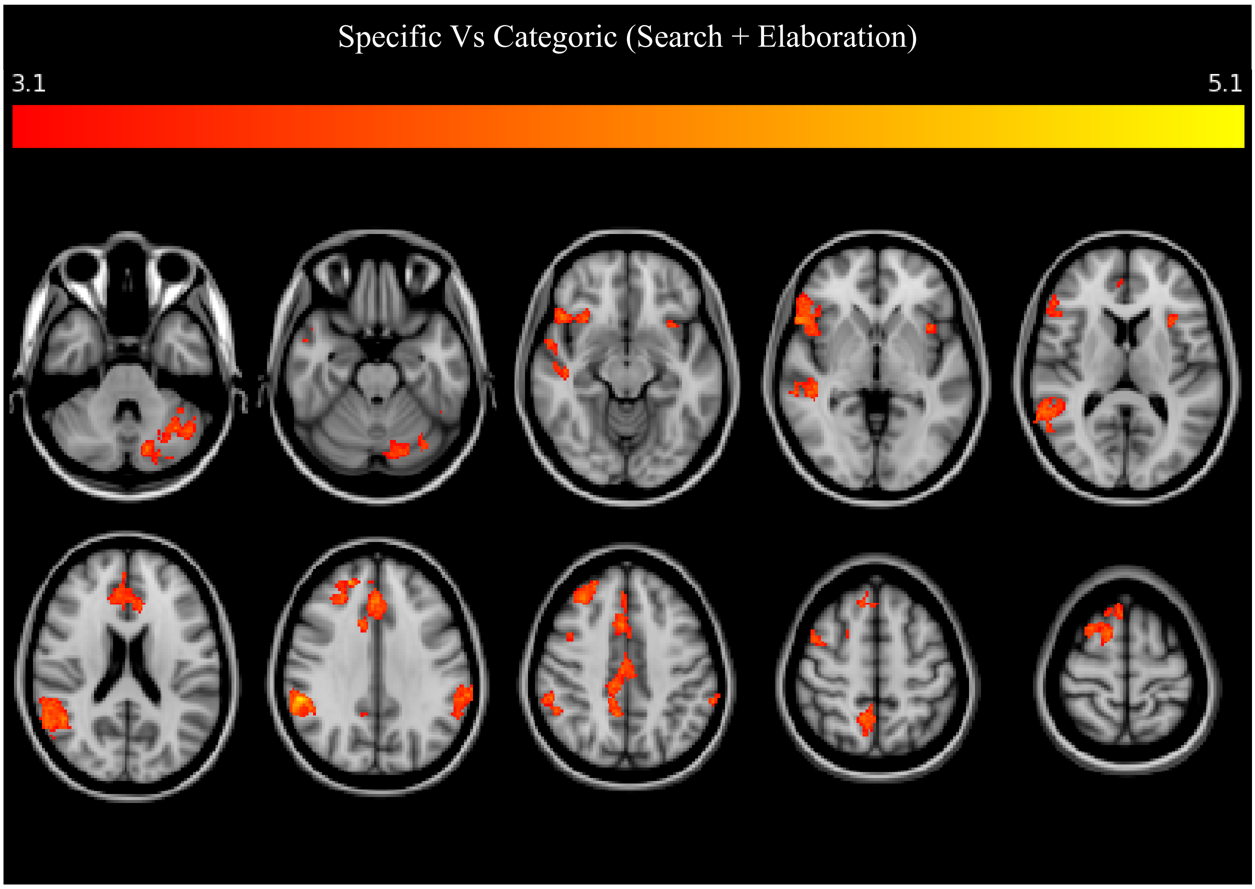


*Figure 1*. Whole-brain mixed-effects analysis of the main effect of memory condition. BOLD activity more strongly associated with specific, compared to categoric retrieval for the whole model (search + elaboration).

| ClusterIndex | Voxels | Z score | MNI X (mm) | MNI Y (mm) | MNI Z (mm) | Atlas Labels |
|---|---|---|---|---|---|---|
| 9 | 2372 | 4.47 | 2 | 22 | 38 | Paracingulate Gyrus (20.1%); Cingulate Gyrus, anterior division (21.1%) |
| 8 | 1595 | 5.16 | 60 | -46 | 26 | Middle Temporal Gyrus, temporooccipital part (12.5%); Supramarginal Gyrus, posterior division (18.7%); Angular Gyrus (27.3%); GM Inferior parietal lobule PF R (11.0%); GM Inferior parietal lobule PFm R (22.3%); GM Inferior parietal lobule Pga R (18.7%) |
| 7 | 1278 | 4.44 | -44 | -56 | -42 | Left VI (15.4%); Left Crus I (50.6%); Left Crus II (21.1%) |
| 6 | 839 | 4.59 | 46 | 22 | -8 | Inferior Frontal Gyrus, pars triangularis (16.8%); Frontal Orbital Cortex (15.1%); GM Broca's area BA45 R (23.3%) |
| 5 | 500 | 4.48 | 20 | 46 | 32 | Frontal Pole (40.5%); Middle Frontal Gyrus (15.7%) |
| 4 | 497 | 4.29 | 48 | -26 | -8 | Superior Temporal Gyrus, posterior division (17.1%); Middle Temporal Gyrus, posterior division (26.4%) |
| 3 | 374 | 4.37 | -62 | -42 | 38 | Supramarginal Gyrus, anterior division (27.8%); Supramarginal Gyrus, posterior division (25.1%); GM Inferior parietal lobule PF L (42.5%); GM Inferior parietal lobule PFm L (18.4%) |
| 2 | 172 | 4.01 | 46 | 4 | 52 | Middle Frontal Gyrus (39.0%); Precentral Gyrus (17.8%); GM Premotor cortex BA6 R (13.7%) |
| 1 | 169 | 3.89 | -32 | 16 | -10 | Insular Cortex (42.4%); Frontal Operculum Cortex (16.9%) |

*Table 1*. Data from analysis of main effects of memory condition, specific > categoric for search and elaboration phases combined, with cluster forming threshold Z>3.1 and (corrected) P < .05.

***Elaboration Model***. Regions which were significantly more active during specific, compared to categoric elaboration are shown in Figure 2 and Table 2**Error! Reference source not found.**. Results from this contrast (specific > categoric) appear to qualitatively overlap with results from the same contrast conducted for the 'whole-block' model. The retrieval of specific (episodic) AMs recorded only for the elaboration phase was associated with greater activity in the middle temporal gyrus, supramarginal gyrus, precuneus, inferior parietal lobe (namely right PGa, bilateral PF and PFm), cerebellum, right Broca's area, and regions in the frontal cortex, including frontal pole and middle frontal gyrus. Of particular interest to the main hypotheses of this chapter, when contrasting specific > categoric BOLD activity, both 'whole' and ‘elaboration’ models revealed greater AG, specifically right PGa, and precuneus activity for specific, compared to categoric memory retrieval.

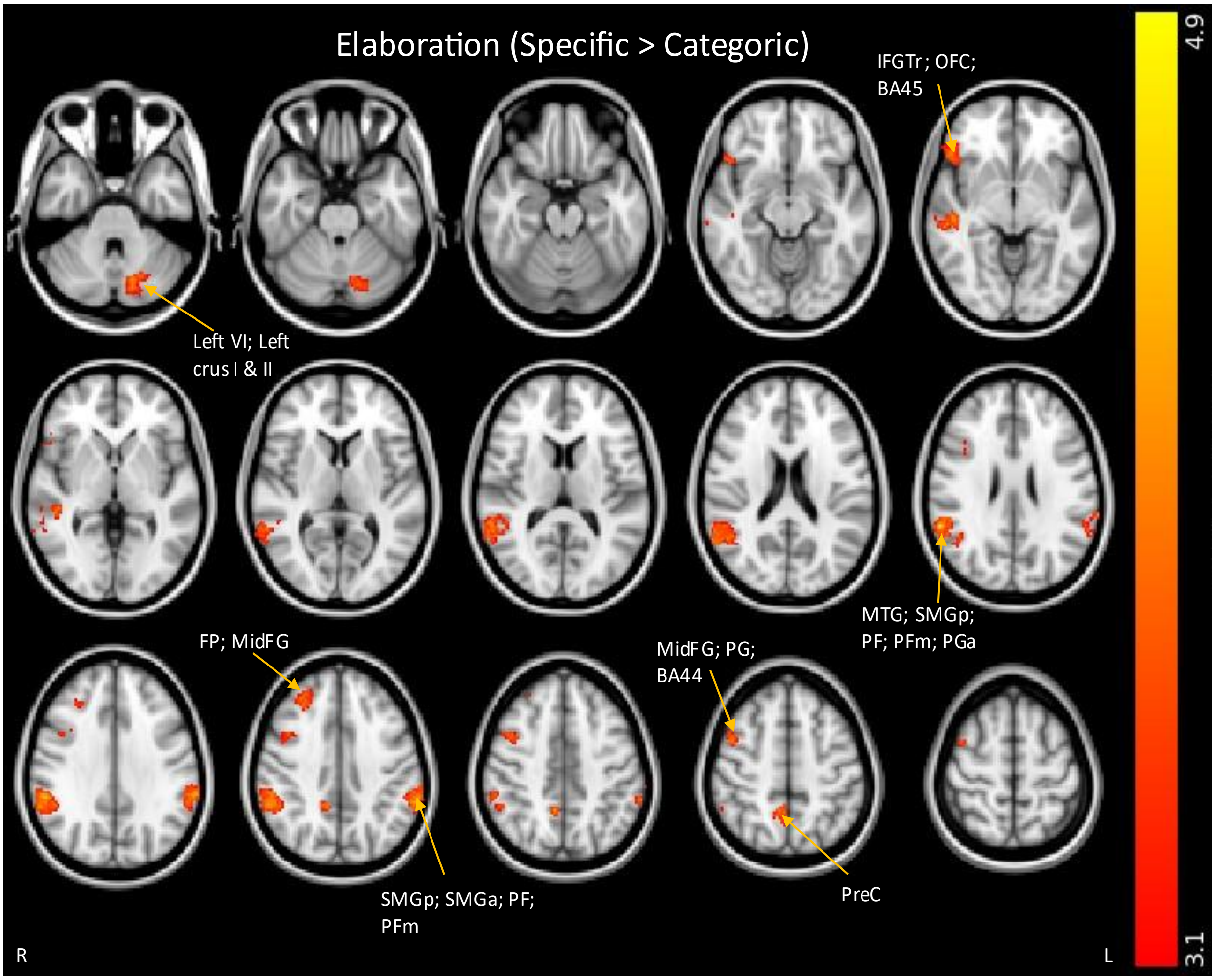


*Figure 2.* Whole-brain mixed-effects analysis of the main effect of memory condition. BOLD activity more strongly associated with specific, compared to categoric retrieval for the elaboration model (elaboration phase only).

| Cluster Index | Voxels | Z score | MNI X (mm) | MNI Y (mm) | MNI Z (mm) | Atlas label(s) |
|---|---|---|---|---|---|---|
| 7 | 1628 | 4.84 | 60 | -46 | 28 | Middle Temporal Gyrus, temporooccipital part (10.5%); Supramarginal Gyrus, posterior division (17.0%); Angular Gyrus (25.5%); GM Inferior parietal lobule PF R (10.0%); GM Inferior parietal lobule PFm R (21.7%); GM Inferior parietal lobule Pga R (17.6%) |
| 6 | 443 | 4.45 | -60 | -42 | 40 | Supramarginal Gyrus, anterior division (27.3%); Supramarginal Gyrus, posterior division (26.2%); GM Inferior parietal lobule PF L (41.8%); GM Inferior parietal lobule PFm L (17.5%) |
| 5 | 372 | 4.39 | -14 | -72 | -30 | Left VI (13.9%); Left Crus I (43.5%); Left Crus II (31.3%) |
| 4 | 234 | 4.03 | 46 | 4 | 52 | Middle Frontal Gyrus (34.2%); Precentral Gyrus (17.9%); GM Broca's area BA44 R (11.1%) |
| 3 | 197 | 4.02 | 46 | 22 | -8 | Inferior Frontal Gyrus, pars triangularis (17.7%); Frontal Orbital Cortex (24.3%); GM Broca's area BA45 R (17.8%) |
| 2 | 179 | 4.04 | 24 | 44 | 40 | Frontal Pole (31.7%); Middle Frontal Gyrus (25.6%) |
| 1 | 169 | 4.17 | 8 | -50 | 48 | Precuneous Cortex (63.8%) |

*Table 2.* Data from analysis of main effects of memory condition, specific > categoric for the elaboration phase.

***Details Model.*** A group analysis of elaboration where the level of details recalled for each trial was included as parametric regressor at the first level, was conducted. This model offers a greater specificity on the memory process by differentiating activity to the standard model using a weighting of 1 regressor and assessing the relationship between activity and the amount of detail being retrieved. Group averages for the parametric (details) contrasts for both specific and categoric elaboration were estimated (Table 3).

| Specific Elaboration | | | | | | |
|---|---|---|---|---|---|---|
| Cluster Index | Voxels | Z score | MNI X (mm) | MNI Y (mm) | MNI Z (mm) | Atlas label(s) |
| 10 | 18795 | 5.35 | -4 | -70 | 24 | Precuneous Cortex (13.9%) |
| 9 | 2354 | 4.66 | 16 | -50 | -48 | Left VI (19.1%); Right VIIIa (10.2%) |
| 8 | 508 | 4.36 | 28 | -60 | -22 | Right VI (58.8%); Right Crus I (25.4%) |
| 7 | 472 | 4.2 | -62 | -42 | 24 | Supramarginal Gyrus, anterior division (18.4%); Supramarginal Gyrus, posterior division (20.9%); Parietal Operculum Cortex (15.0%); GM Inferior parietal lobule PF L (43.8%); GM Inferior parietal lobule PFcm L (20.4%); GM Inferior parietal lobule PFm L (15.0%) |
| 6 | 463 | 4.15 | -44 | -78 | 16 | Lateral Occipital Cortex, superior division (59.0%); GM Inferior parietal lobule PGp L (43.3%) |
| 5 | 345 | 4.42 | 56 | -44 | 22 | Middle Temporal Gyrus, temporooccipital part (10.7%); Supramarginal Gyrus, posterior division (24.5%); Angular Gyrus (22.2%); GM Inferior parietal lobule PFm R (18.4%); GM Inferior parietal lobule Pga R (15.2%) |
| 4 | 272 | 4.04 | 40 | 10 | 8 | Insular Cortex (20.3%); Central Opercular Cortex (19.4%) |
| 3 | 222 | 4.04 | -30 | 46 | 24 | Frontal Pole (57.9%) |
| 2 | 204 | 4.44 | 50 | -4 | -18 | Temporal Pole (12.6%); Superior Temporal Gyrus, anterior division (14.2%); Middle Temporal Gyrus, anterior division (10.1%) |
| 1 | 164 | 3.97 | 60 | -28 | 38 | Supramarginal Gyrus, anterior division (24.0%); Parietal Operculum Cortex (24.9%); GM Inferior parietal lobule PF R (24.6%); GM Inferior parietal lobule PFcm R (29.8%); GM Inferior parietal lobule PFop R (15.5%); GM Secondary somatosensory cortex / Parietal operculum OP1 R (22.8%) |
| Categoric Elaboration | | | | | | |
| 1 | 192 | 3.91 | -14 | 20 | 4 | Left Caudate (50.4%); Left Putamen (10.8%) |

*Table 3.* Details Model (elaboration phase). Whole-brain BOLD activity for categoric and episodic elaboration with details as a regressor.

We also assessed the main effect of memory type, by contrasting activity for specific greater that categoric retrieval and vice versa, for the parametric contrast (details). Table 4 demonstrates the significantly stronger positive association between activity recorded during specific elaboration and self-reported details, compared to categoric elaboration. These regions included the precuneus cortex, cuneal cortex, subregions of the superior parietal lobule (right 7A, 7M and 7P), as well as subregions of the cerebellum (Figure 3). No clusters with Z score higher than 3.1 were found for the contrast categoric > specific.

| Cluster Index | Voxels | Z score | MNI X (mm) | MNI Y (mm) | MNI Z (mm) | Atlas label(s) |
|---|---|---|---|---|---|---|
| 3 | 459 | 4.14 | 18 | -64 | 28 | Precuneous Cortex (37.5%); Cuneal Cortex (18.1%) |
| 2 | 154 | 3.86 | 6 | -64 | 62 | Precuneous Cortex (59.0%); GM Superior parietal lobule 7A R (22.0%); 7M R (11.1%); 7P R (21.4%) |
| 1 | 151 | 4.04 | -24 | -42 | -44 | Left VIIIa (16.4%); Left VIIIb (12.4%); Left X (14.9%) |

*Table 4.* BOLD activity for specific, compared to categoric memory retrieval, with details as a regressor. Clusters were determined by Z<3.1, and a corrected cluster significance threshold of P = 0.05. For each region of activation, the coordinates of the peak within each structure is reported, as indicated by the highest Z-score.

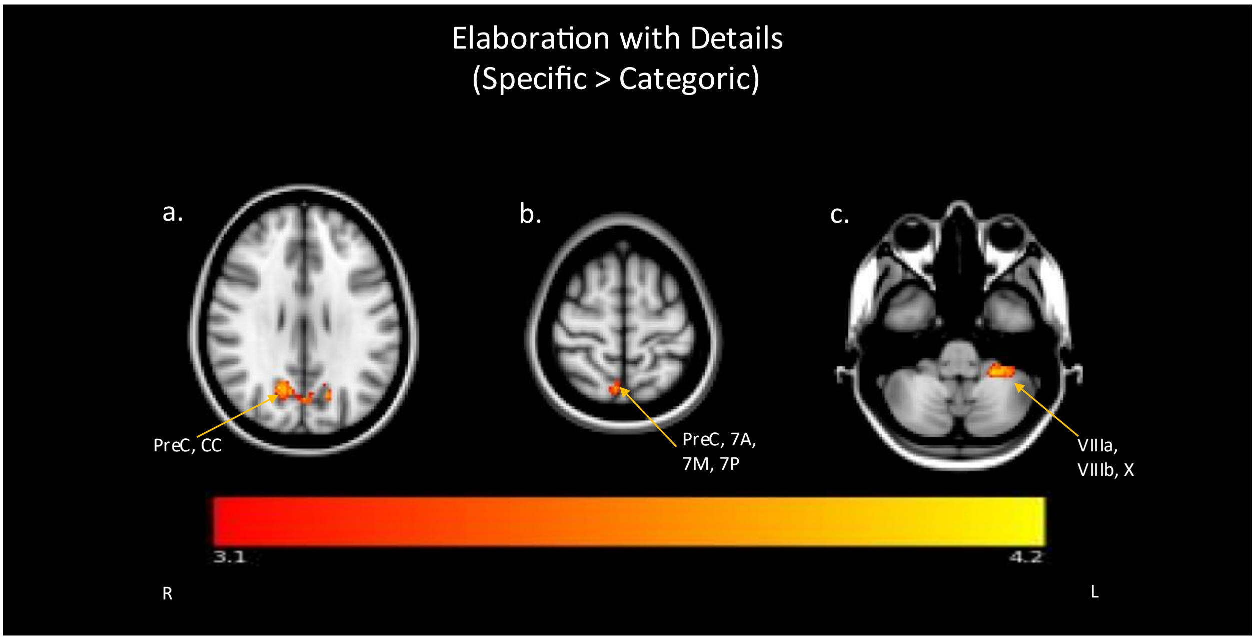


*Figure 3.* BOLD activity associated with specific elaboration (with details as regressor), compared to categoric elaboration. Peak co-ordinates are reported, a. = praecuneus and cuneal cortex (CC). b. = praecuneus, right superior parietal lobule (7A, 7M and 7P); c. = left VIIIa, left VIIIb and left X in the cerebellum.

### Masked Analyses

To investigate whether the AG is sensitive to the richness of the content that is being retrieved during elaboration and assess different involvement of AG subregions PGa and PGp. We also had prior hypotheses of a positive relationship between AG and details, irrespective of whether the memory was episodic or semantic in nature. We first assessed the group relationship between activity and number of details retrieved for each memory type separately. For completeness we also contrasted specific Vs categoric conditions to assess differences of the relationship between activity and details between memory types. To assess activity in the AG masks were created for each subregion separately (left and right PGa and PGp). These were then combined to provide a more general overview of the role of the AG in rich memory retrieval and create a left and right AG mask. In line with extensive evidence demonstrating key involvement of both the hippocampus and precuneus in the retrieval of AM, we also included masks for these two brain areas (left and right hippocampus and precuneus).

#### *Differences Between Memory Types: Specific Vs Categoric*

When assessing whether activity is differentially recruited for specific, compared to categoric memories, no differentiation was seen for the categoric > specific contrast. Greater BOLD activity recorded during specific elaboration, compared to categoric elaboration was seen for right AG (including PGa and PGp) and precuneus (Table 5).

| Contrast | Mask | Voxels | P value | MNI (mm) | XMNI (mm) | Y MNI Z (mm) |
|---|---|---|---|---|---|---|
| Categoric > Specific | No masks reached significance | | | | | |
| Specific > Categoric | Left AG | - | - | - | - | - |
| | Right AG | 508 | 0.004 | 58 | -52 | 16 |
| | Left Pga | - | - | - | - | - |
| | Right Pga | 426 | 0.002 | 58 | -52 | 16 |

| | | | | | |
|---|---|---|---|---|---|
| Left PGp | - | - | - | - | - |
| Right PGp | 31 | 0.025 | 52 | -70 | 20 |
| Right PGp | 31 | 0.029 | 50 | -60 | 20 |
| Left HP | - | - | - | - | - |
| Right HP | - | - | - | - | - |
| Precuneus | 24 | 0.038 | 8 | -48 | 42 |

*Table 5.* Main effects of comparing memory conditions during elaboration. Significant results only seen for the specific > categoric masked analyses contrasts.

***Memory Retrieval and Details during Elaboration phase***

Looking more specifically at activity in the selected masks to test our hypothesis of whether the AG is sensitive to richness of retrieved content, activity for the parametric regressor (details) was averaged across the group. Again, as seen for the whole-brain group 'Elaboration' analysis, ratings were assessed in conjunction with activity in the elaboration phase of the task. Results from non-parametric analysis can be seen in Table 6 and Figure 4. The analysis included the following contrasts: categoric elaboration with details as regressor, specific elaboration with details as regressor. For categoric AMs, a significant positive relationship with number of details was observed between BOLD activity in the left AG, specifically left PGp, and bilateral hippocampus. For specific AMs, a significant positive relationship to level of details retrieved was seen for bilateral AG, specifically, bilateral PGp activity and right PGa activity, as well as in bilateral hippocampus and precuneus.

| Contrast | Mask | Voxels | P value | MNI X (mm) | MNI Y (mm) | MNI Z (mm) |
|---|---|---|---|---|---|---|
| Categoric Details | Left AG | 57 | 0.024 * | -48 | -70 | 20 |
| | Left AG | 31 | 0.028 * | -32 | -88 | 34 |
| | Right AG | - | - | - | - | - |
| | Left Pga | - | - | - | - | - |
| | Right Pga | - | - | - | - | - |
| | Left PGp | 204 | 0.015 * | -48 | -70 | 20 |
| | Right PGp | - | - | - | - | - |
| | Left HP | 128 | 0.01 * | -32 | -20 | -16 |
| | Right HP | 71 | 0.014 * | 28 | -18 | -14 |
| | Precuneus | - | - | - | - | - |
| Specific Details | Left AG | 414 | 0.006 * | -36 | -84 | 28 |
| | Right AG | 884 | 0.001 * | 46 | -80 | 18 |
| | Left PGa | - | - | - | - | - |
| | Right PGa | 227 | 0.012 * | 48 | -48 | 26 |
| | Left PGp | 451 | 0.004 * | -36 | -86 | 26 |
| | Right PGp | 685 | 0.001 * | 44 | -82 | 20 |
| | Left HP | 416 | < 0.001 * | -34 | -34 | -8 |
| | Right HP | 355 | 0.001 * | 32 | -14 | -18 |
| | Precuneus | 3615 | < 0.001 * | 18 | -58 | 12 |

*Table 6.* Main effects of memory elaboration for specific and categoric memories, with details as regressor.* represents agreement of significant results between parametric and non-parametric masked analyses.

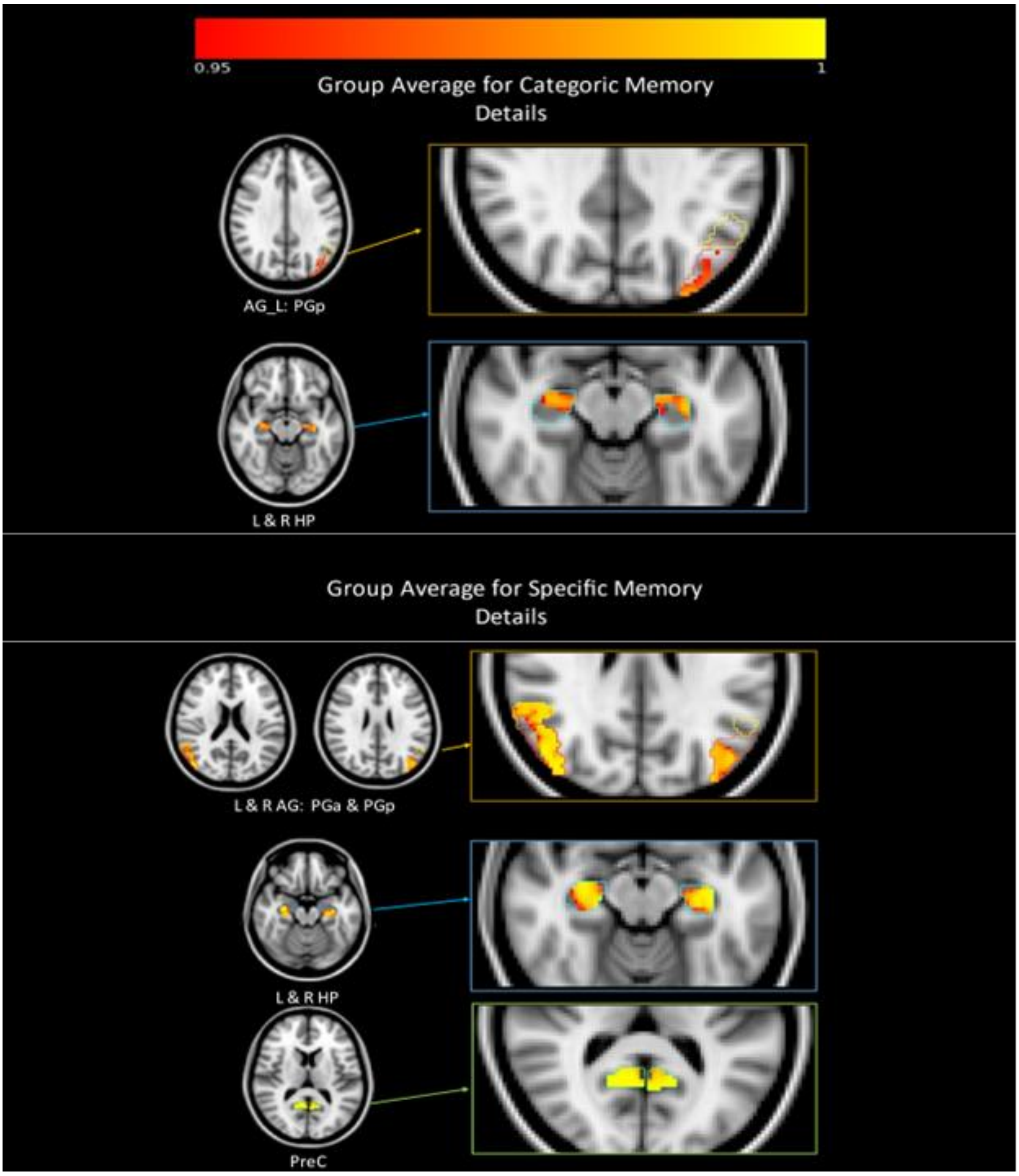


*Figure 4*. Main effect of categoric and specific retrieval, with details as regressor (TFCE corrected $p$ < .05, p-value colour bar shown). Mask outline is also presented, with AG mask subdivided into PGa and PGp. Non-parametric masked analyses reached significance for bilateral AG (left PGp, right PGa & PGp), bilateral hippocampus and precuneus for the specific condition. Significant results only seen for left AG (specifically PGp) and bilateral hippocampus for the categoric condition.

Finally, when assessing whether activity based on level of details recalled is differentially recruited for specific, compared to categoric memories, no differentiation was seen for the categoric > specific contrast. A stronger positive relationship between the BOLD activity recorded during specific elaboration and details was seen compared to the equivalent contrast for categoric elaboration for the right AG (including PGa and PGp), left hippocampus and precuneus. For comparison, at the whole-brain level, only the precuneus and left cerebellum were identified as being more positively associated with details in the comparison specific > categoric (see Figure 5 and Table 7).

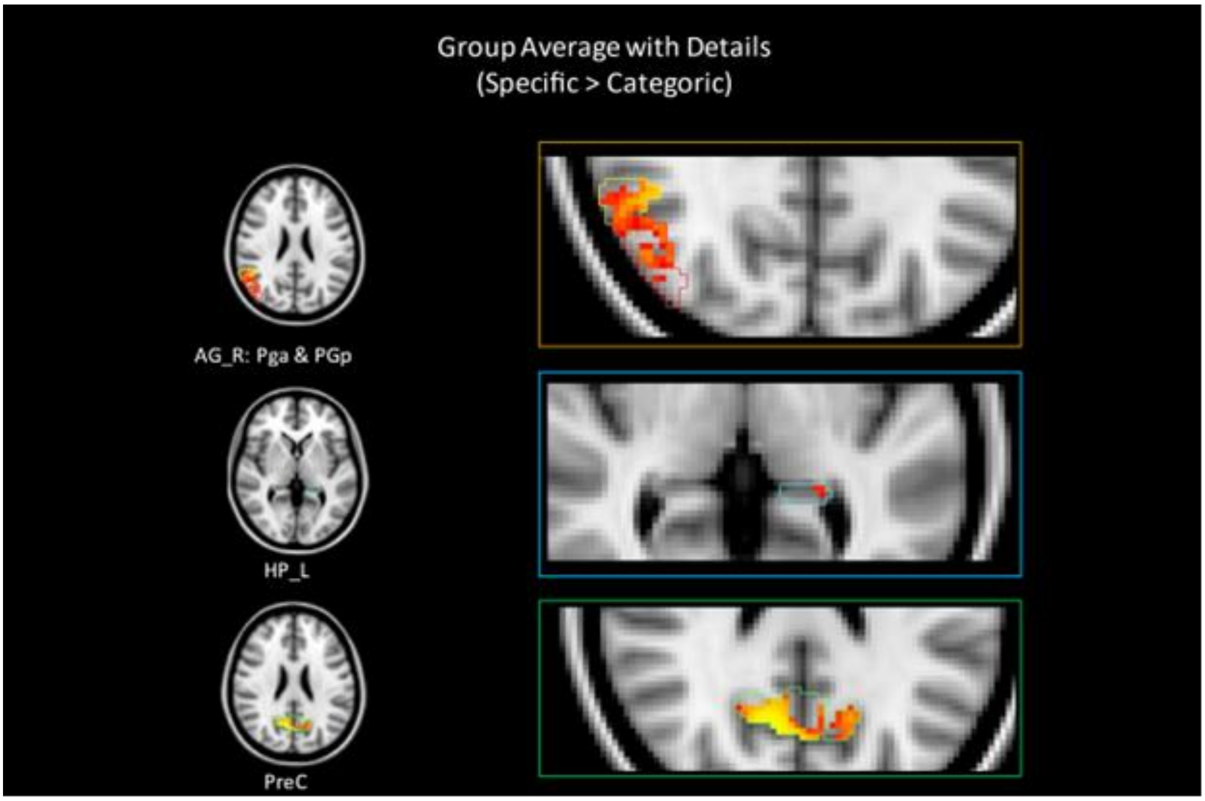


*Figure 5*. Main effect of memory condition (specific > categoric), with details as regressor (TFCE corrected $p < .05$, p-value colour bar shown). Mask outline is also presented, with AG mask subdivided into PGa and PGp. Non-parametric masked analyses reached significance for right AG (PGa and PGp), left hippocampus and precuneus.

| Contrast | Mask | Voxels | P value | MNI<br>(mm) | XMNI<br>(mm) | Y<br>MNI Z (mm) |
|---|---|---|---|---|---|---|
| Categoric > Specific | No masks reached significance | | | | | |
| Specific > Categoric | Left AG | - | - | - | - | - |
| | Right AG | 392 | 0.012 | 48 | -50 | 24 |
| | Left Pga | - | - | - | - | - |
| | Right Pga | 199 | 0.01 * | 48 | -50 | 24 |
| | Left PGp | - | - | - | - | - |
| | Right PGp | 187 | 0.01 * | 42 | -68 | 28 |
| | Left HP | 6 | 0.041 * | -24 | -38 | 2 |
| | Right HP | - | - | - | - | - |
| | Precuneus | 1987 | 0.002 * | 10 | -66 | 24 |

*Table 7*. Main effects of comparing memory conditions, with details as regressor. Significant results only seen for the specific > categoric masked analyses contrasts.* represents agreement of significant results between parametric and non-parametric masked analyses.
